# Long-Stokes-Shift mScarlet3 as a Structural Marker for Two-Photon Imaging

**DOI:** 10.64898/2026.04.12.718060

**Authors:** Sangyu Xu, Xianyuan Zhang, King Yee Cheung, Yishan Mai, Yue Wu, Adam Claridge-Chang

## Abstract

Two-photon imaging with genetically encoded sensors is widely used to monitor neurophysiology. An additional fluorescent protein can provide anatomical landmarks for cell-type identification and motion detection. However, monomeric red fluorescent proteins often require a dedicated excitation laser. We made transgenic *Drosophila* with a long-Stokes-shift mScarlet variant (*LSSmScarlet3*) to image alongside green sensors with a single 920-nm laser. We describe excitation and emission spectra of the expressed protein and show robust fluorescence at 920 nm *in vitro* and *in vivo* with minimal channel crosstalk. This approach can reduce equipment complexity and cost while placing functional calcium dynamics in their anatomical context.

## Description

Understanding neural circuits requires monitoring both neural activity and anatomical relationships. Genetically encoded calcium indicators based on blue-excitable **g**reen **f**luorescent **p**rotein (GFP) are widely used, for example, the **G**FP–**Ca**l**m**odulin–M13-**p**eptide (*GCaMP*) series of calcium sensors. Such sensors enable cellular-resolution functional imaging, but calcium signals are transient and can afford limited structural context. Alongside sensors, researchers sometimes co-express **r**ed **f**luorescent **p**roteins (RFPs) as structural markers (e.g., tdTomato or mCherry) to provide landmarks for cellular identification and sample-motion detection. Single-laser excitation of red fluorescent proteins away from their excitation maxima (∼920 nm rather than at their ∼1040 nm peak) has been used for dual-color structural-functional imaging (Tillo et al. 2010; Ozbay et al. 2018), but relies on the tandem dimer tdTomato, whose two-photon cross-section at this wavelength is an order of magnitude higher than that of monomeric RFPs such as mCherry (∼35–40 GM vs. ∼5 GM) (Drobizhev et al. 2011). The resulting doubling of molecular weight complicates protein-fusion functionality (Shaner et al. 2005). Alternatively, a dedicated 1040 nm laser can be added to excite monomeric RFPs at their peak (Gee et al. 2014; Bugeon et al. 2022), resulting in multi-laser setups that increase equipment cost, system complexity, and phototoxicity. A monomeric red fluorophore with robust excitation at ∼920 nm could therefore retain the simplicity of single-wavelength imaging while avoiding a tandem dimer, improving signal levels at low expression, and broadening applications to include fusion proteins and subcellular markers.

Long-Stokes-shift (LSS) fluorescent dyes exhibit large separations between excitation and emission peaks (Shcherbakova et al. 2012). Meanwhile, there has been considerable effort in developing LSS fluorescent proteins through diverse approaches, including excited-state proton transfer, chromophore modifications, and protein engineering (Piatkevich et al. 2010; Santos et al. 2021). Originally developed to improve FLIM-FRET (fluorescence lifetime imaging microscopy-Förster resonance energy transfer) sensors by reducing spectral overlap between donor and acceptor fluorophores, LSS variants have found broader applications in multicolor imaging (Shcherbakova et al. 2012; Piatkevich et al. 2010). A recently developed LSS variant of mScarlet, *LSSmScarlet3*, exhibits a 128 nm Stokes shift (ex 466 → em 598 nm) and, compared to previous versions, has improved pH stability and photostability (Subach et al. 2022). Similarly, **m**onomeric **cy**an-excitable **RFP 3**, *mCyRFP3 (Kim et al. 2022)*, has been reported as a functional fusion tag in cell-based imaging and has recently been incorporated into Drosophila transgenic reagents (FlyBase 2025). Here, as a complementary contribution, we describe a genetically encoded *LSSmScarlet3* as a structural marker for single-laser two-color 2P imaging, demonstrating efficient excitation at 920 nm. Alongside *GFP*-based sensors, *LSSmScarlet3* provides a simplified approach for dual-channel functional–structural imaging.

To verify whether a long-Stokes-shift RFP can be imaged with a 920 nm laser in *Drosophila* brains, we synthesized a *Drosophila* codon-optimized version of LSSmScarlet3. In fixed *Drosophila* tissue, *LSSmScarlet3* was well expressed and readily distributed to the lobes of the mushroom body, indicating good neuronal expression and distribution (**Figure 1A**). On a spectrally tunable multiphoton microscope, *LSSmScarlet3* had a peak excitation at 940 nm, with two smaller local maxima at 860 nm and 1040 nm (**Figure 1B**). At a frequently used 2P excitation wavelength of 920 nm, the mean ROI intensity measured was 99 arbitrary units (arb. u.), which was around 75% of the 940-nm peak intensity (132 arb. u.). Emission curves for the three excitation wavelengths tested (**Figure 1C**) show a peak emission wavelength around 620 nm.

**Figure 1.**
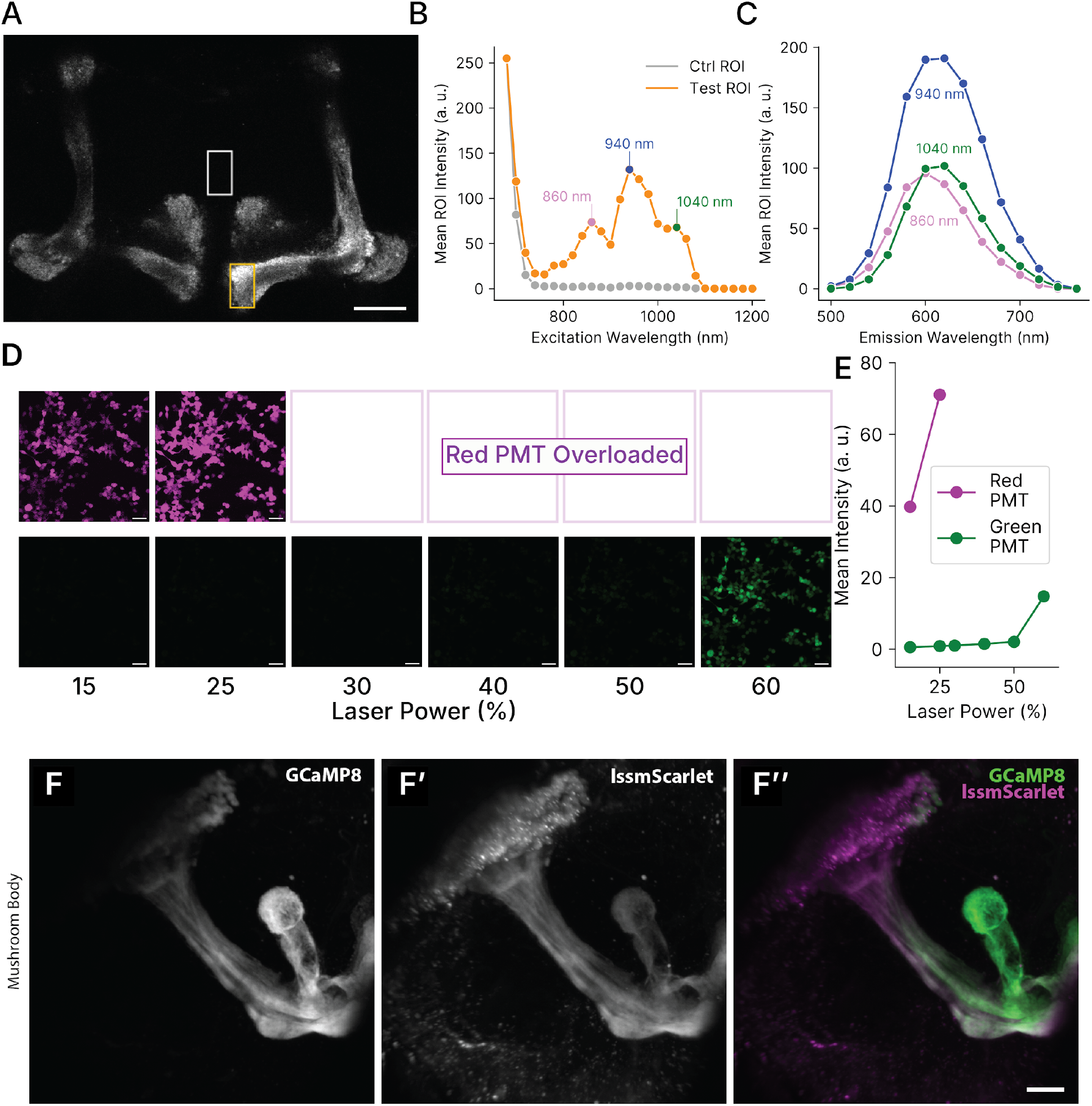
Two-photon imaging of a *Drosophila* codon-optimized *LSSmScarlet3*. **A)** An image of the mushroom body expressing *LSSmScarlet3* in a fixed *MB247-Gal4>UAS-LSSmScarlet3* fly collected with 940 nm excitation and 620 nm collection. White box: control ROI without *LSSmScarlet3* expression. Orange box: test ROI with *LSSmScarlet3* expression. Scale bar: 20 µm. **B)** Excitation spectrum of *LSSmScarlet3* in the sample shown in **A**. Three local maxima were further tested for emission: 860 nm, 940 nm, and 1040 nm. **C)** Emission spectra of excitation at 860 nm, 940 nm, and 1040 nm. **D)** *LSSmScarlet3* expressed in live HEK293T cells under the control of a CMV promoter excited at various 920 nm laser power settings and images collected with a green and a red PMT. At laser power above 25 %, the red PMT was overloaded by the light emitted. Green signals began to emerge at above 50 % laser power. **E)** Quantification of mean intensity of each image in **D** as a function of laser power. **F)** Maximum intensity projection image of the mushroom body calyx and α-lobe in a live *MB247>GCaMP8m;LSSmScarlet3 Drosophila* brain. Both fluorophores were simultaneously excited by the 920 nm laser at 20% power. (F) GCaMP8m signals detected with the green-filtered PMT at 80% gain, (F′) LSSmScarlet3 signals detected with the red-filtered PMT at 70% gain, and (F′′) merged signals. Scale bar = 20 µm.

Having established the spectral properties of the fluorophore, we wanted to measure its performance on a standard 2P microscope equipped with one 920 nm laser and two photomultiplier tubes (PMTs) for detection: one green, one red (**Figure 1D**). When expressed in **h**uman **e**mbryonic **k**idney **293T** (HEK293T) cells under the control of a CMV promoter, and with 920 nm excitation, *LSSmScarlet3* was detected with the red PMT. As we varied the laser power input, we observed robust detection of *LSSmScarlet3* by the red PMT at low gain, whereas the green PMT did not detect any signal below 50 % laser power. Above 25 % laser power, the red PMT was overloaded. Only at around 60 % laser power did we observe substantial cross-talk signal on the green PMT (**Figure 1D, E**). This separation between RFP and GFP-channel detection indicates that, with appropriate filtering, *LSSmScarlet3* is suitable for marking cells without interference with the green channel.

Co-expression of *LSSmScarlet3* and *GCaMP8m* in the mushroom body under UAS control, driven by MB247-Gal4, showed robust co-detection (**Figure 1F**) *in vivo*.

We demonstrate the utility of *LSSmScarlet3* as a structural marker alongside *GCaMP8m* using a single laser for excitation. This arrangement promises substantial advantages over existing approaches, including simplicity in setup, elimination of sequential scanning for potential reduction of phototoxicity, and better temporal registration. Strong excitation at 920 nm is ideal for most existing 2P setups, and we validated that the long Stokes shift minimizes crosstalk between green and red channels. Compared to other long Stokes shift FPs currently available, our transgenic fly is validated and ready to use.

*LSSmScarlet3* is quite bright in our hands. Given the low threshold for PMT saturation from the highly expressed *LSSmScarlet3* in HEK293T cells **(Figure 1D)**, careful titration of expression levels may be needed. On the other hand, this also reassures users of highly detectable signals when expression levels are low, for example, in a fusion protein. This opens up many possibilities, such as co-imaging neuronal activity with synaptic markers, tagging proteins with subcellular localization signals, or monitoring reference transcription levels.

Transgenic *LSSmScarlet3* provides a practical solution for dual-channel 2P imaging in *Drosophila*. Single-laser, two-color imaging opens up avenues for developing *LSSmScarlet3* variants for specific biological markers. For applications with strong promoters where structural bulk is not a concern, using tdTomato excited with ∼920 nm remains a viable option. LSSmScarlet3 offers particular advantages when a monomeric tag is needed, such as with fusion proteins, weak promoters, or subcellular localization signals where expression levels may be limiting.

## Methods

### Fly stocks and husbandry

*Drosophila* cultures were maintained on standard cornmeal-based medium (1.25 % w/v agar, 10.5 % w/v dextrose, 10.5 % w/v maize, and 2.1 % w/v yeast) (Temasek Life Sciences Laboratories 2018). Stocks were kept at room temperature unless otherwise specified. Crosses were performed at 25 ºC in a day-night cycling incubator (Mir-253, Sanyo). Adult male and female flies, aged 5–10 days, were used for experiments. UAS-jGCaMP8m (Zhang et al. 2023) was obtained from Bloomington Drosophila Stock Center (BDSC #92590). The MB247-Gal4 stock (Zars et al. 2000) was provided by Dr Hiromu Tanimoto (Tohoku University, Japan).

### Construct generation

*LSSmScarlet3* coding sequence was codon-optimized for *Drosophila* by GenScript (Piscataway, NJ, USA) and cloned into *pcDNA3*.*4* (Thermo Fisher Scientific, Waltham, MA, USA; cat. no. V880-01) to generate the mammalian expression construct *pcDNA-DmLSSmScarlet3* and into *pJFRC81-10XUAS-IVS-Syn21-GFP-p10* vector (Pfeiffer et al. 2012) between XhoI and XbaI restriction sites to generate *Drosophila* expression construct *UAS-DmLSSmScarlet3*, which was then injected into *yw* flies at the attp40 site balanced against Curly of Oster by BestGene (La Jolla, CA, USA). The *pJFRC81-10XUAS-IVS-Syn21-GFP-p10* plasmid was a gift from Gerald Rubin (Addgene plasmid #36432; http://n2t.net/addgene:36432;RRID:Addgene_36432)

### Excitation and emission scanning of mushroom body tissue

Fly brains expressing *LSSmScarlet3* were dissected in cold PBS and fixed in 4 % PFA for 30 min and then mounted in PBS in 2 stacked reinforcement rings (Suremark) pasted on 1.0–1.2 mm-thick 25 × 75 mm glass slides (Biomedia), and then sealed with 170 µm-thick 18 × 18 mm glass cover slips with clear nail polish. Samples were imaged on a Leica TCS SP8 DIVE multiphoton microscope (Leica Microsystems, Wetzlar, Germany) as previously described (Cheung et al. 2026) with the exception that in the emission spectrum measurements the excitation wavelengths were fixed at 860 nm, 940 nm, or 1040 nm. Image acquisition and spectral detection were controlled using Leica LAS X software. Mean intensity of ROIs in images was calculated using Fiji (Schindelin et al. 2012). Excitation and emission spectra graphs were generated using matplotlib in Python.

### Cell culture and transfection

HEK293T cells were seeded into small tissue culture dishes to ∼ 70 % confluence in Dulbecco's Modified Eagle Medium (Thermo Fisher Scientific, Waltham, MA, USA) containing 10% Fetal Bovine Serum (Thermo Fisher Scientific, Waltham, MA, USA) and transfected with *pcDNA3*.*4*-*DmLSSmScarlet3* constructs using Lipofectamine 3000 (Thermo Fisher Scientific, Waltham, MA, USA) in Opti-MEM™ Medium (Thermo Fisher Scientific, Waltham, MA, USA) according to a standard protocol (Thermo Fisher Scientific 2016) in 35 mm cell culture dishes (Nunc, catalog #153066; Thermo Fisher Scientific, Waltham, MA, USA). The cells were imaged 48 hours after transfection.

### 2-photon imaging of *LSSmScarlet3* in HEK293T cells

Cell culture expressing *LSSmScarlet3* was imaged with A-SCOPE (Thorlabs, NJ, USA) as previously described (Cheung et al. 2026). Laser power was set to 30 mW. Green PMT was bandpass filtered between 500 nm and 550 nm, and the red PMT was bandpass filtered between 570 nm and 640 nm. Detector gains were set to 5%.

### Simultaneous 2-photon imaging of *GCaMP8m* and *LSSmScarlet3* in *Drosophila* mushroom body

Flies were mounted in a custom 3D-printed chamber using candle wax. Fly heads were opened using a pair of sharp Dumont No. 5 forceps (Fine Science Tools) and microdissection scissors (Fine Science Tools). A drop of PBS was then placed on the open fly head to keep the brain moist and to act as the objective immersion medium.

*Z*-stack images were acquired on a FEMTO3D Atlas Plug & Play 2-photon microscope (Femtonics, Budapest, Hungary) equipped with a 16× objective lens. Laser power of the 920 nm laser was set to 20% (112 mW). *GCaMP8* emission light was collected by a green-filtered PMT at 80% gain, while *LSSmScarlet3* emission light was collected by a red-filtered PMT at 70% gain.

### Image processing

The *Drosophila* mushroom body 2-photon image stack was subjected to Gaussian blur 3D processing (X: 0, Y: 0, Z: 2) and maximum intensity projection processing in Fiji (Schindelin et al. 2012). For the purpose of visualizing dual-channel images, signal collected by the green-filtered PMT was pseudo-colored to green, while signal collected by the red-filtered PMT was pseudo-colored to magenta.

### Data analysis and visualization

Preprocessed data from the Fiji workflow were analyzed and visualized using custom Jupyter Notebooks running Python (version 3.10.16) with pandas (version 2.1.4), NumPy (version 1.26.4), and seaborn (version 0.12.2).

## Data and Code Availability

Data and code used to generate the figure in this paper, as well as construct sequences and maps, are available as a Zenodo repository (DOI: 10.5281/zenodo.19493335).

## Author Contributions

S. X.: conceptualization, funding acquisition, methodology, investigation, data curation, formal analysis, visualization, writing — original draft, writing — reviewing and editing

X. Z.: methodology, investigation, writing — reviewing and editing

K. Y. C.: methodology, investigation, data curation, formal analysis, validation, writing — reviewing and editing

Y. M.: methodology, investigation, data curation, validation, writing — reviewing and editing

Y. W.: methodology, investigation, writing — reviewing and editing

A. C.-C.: conceptualization, supervision, funding acquisition, writing — reviewing and editing

## Acknowledgements

We thank Masahiro Fukuda, Hyunsoo Shawn Je (Programme in Neuroscience and Behavioural Disorders, Duke-NUS Medical School) for insightful and productive discussions, Danesha Suresh and Zhiyi Zhang for technical and administrative assistance, and Dénes Pálfi and Viktória Kiss (Femtonics HQ) for assistance with two-photon imaging on FEMTO3D Atlas Plug & Play. We additionally thank Dr. Michael Hausser (University of Hong Kong) and Dr. Jung Kim (University of Hong Kong) for providing temporary access to the FEMTO3D Atlas and lab space. The design of the custom chamber used for imaging was kindly provided by Dr. Anissa Kempf (University of Basel). We thank BDSC and Dr. Hiromu Tanimoto for providing fly stocks.

## Funder Information Declared

National Medical Research Council (https://ror.org/04x3cxs03), MOH-OFYIRG20nov-0051, MOH-001397-01

Ministry of Education, Singapore (https://ror.org/01kcva023), MOE-T2EP30222-0018, MOE-T2EP30223-0009, FY2022-MOET1-0001

Duke-NUS Medical School (https://ror.org/02j1m6098)

